# Targeting Glioblastoma Cell State Plasticity for Enhanced Therapeutic Efficacy

**DOI:** 10.1101/2025.09.08.674897

**Authors:** Stefano M. Cirigliano, Richa Singhania, James Nicholson, Isha Monga, Yushan Wan, Caroline Haywood, Ashlesha Muley, Skylar Giacobetti, Howard A. Fine

**Affiliations:** Department of Neurology, Meyer Cancer Center, Weill Cornell Medicine, New York, NY, 10021 USA

**Keywords:** tumor neurobiology, glioblastoma, single-cell, treatment resistance, cancer stem cell, cerebral organoids

## Abstract

Glioblastoma (GBM) is the most common and deadly primary brain cancer, with limited therapeutic options. Treatment failure has been associated with intratumoral heterogeneity and the acquisition of a pronounced mesenchymal-like (MES-L) phenotype after recurrence. Here, we have screened a panel of drugs with diverse mechanisms of action across two patient-derived glioblastoma stem cells (GSCs) to characterize the dynamics of drug-mediated transcriptomic cellular state changes. Our results demonstrate that anti-tumor drugs induce significant but reversible alterations in cellular state distribution at the single-cell level in a drug-specific manner, influencing transitions between mesenchymal and the neurodevelopmental astrocytic-like (AC-L) states. Utilizing barcoded analysis in our recently developed ex vivo glioblastoma cerebral organoid (GLICO) model, we discerned distinct cell state sensitivities to the MES-L enhancing histone deacetylase inhibitor, panobinostat, which are contingent on the inducible modulation of the mesenchymal transcription factor FOSL1. The strategic combination of MES-L enhancing and MES-l suppressing genetic perturbations or drugs significantly increases anti-glioma activity in a strategy we call state-selective lethality. Overall, our findings highlight the critical role of cell state plasticity in the response of GSCs to anti-tumor therapeutic stress and underscore the potential for novel GBM combination drug strategies.

## Introduction

Glioblastoma multiforme (GBM) is an aggressive and highly infiltrative primary brain tumor characterized by treatment resistance, high recurrence rate, and low survival. likely reason that GBMs are so resistant to effective treatment is their extensive intratumoral heterogeneity and plasticity across a spectrum of environmental and stress-associated transcriptional pseudo-hierarchical cellular states. Although variously named and defined, the most commonly recognized cellular states are the astrocyte (AC)-like, mesenchymal (MES)-like, oligodendrocyte progenitor cell (OPC)-like, and neural progenitor cell (NPC)-like states (Neftel et al., 2019). Spatially resolved sequencing of whole GBM tumors also showed region-specific genetic subclones and microenvironment-driven expression programs in the tumor core versus invasive edges. Within a single tumor, glioma stem cells (GSCs) tend to occupy different niches (e.g. MES-L GSCs often in immune-reactive, macrophage-rich regions) and collectively drive tumor growth and recurrence (Greenwald et al., 2024). This cellular state diversity and plasticity may allow the tumor to adapt and resist therapeutic stress such as radiation and chemotherapy, therefore contributing to the poor prognosis associated with GBM – a hypothesis yet to be tested in detail in preclinical relevant models.

GBMs employ several mechanisms of therapeutic stress tolerance, including entering quiescence to evade drug targeting (Antonica et al., 2022) and reprogramming metabolism under hypoxic conditions (Singh et al., 2025). Epigenomically mediated cell state plasticity enables GBM cells to transition between states in response to environmental pressures and may therefore represent another powerful mechanism of treatment resistance, consistent with the observation that therapeutic stress can induce a proneural-to-mesenchymal shift, resulting in a more aggressive, drug-resistant phenotype (Park et al., 2024). GBM cells with greater cell state plasticity may thus be more resistant to chemotherapy and radiotherapy. Indeed, combinatorial treatments aimed at blocking key plasticity factors such as FOSL1 have demonstrated synergistic efficacy in preclinical models (Chen et al., 2022). If treatment-specific “refuge cellular states” are less sensitive to certain cytotoxic stresses, then targeting GBM cell states and their plasticity offers a promising therapeutic strategy to overcome the treatment resistance commonly associated with these tumors. By designing novel therapeutic strategies focused on the dynamic nature of these cellular states, it may be possible to disrupt tumor’s adaptive mechanisms and improve patient outcomes, with the advantage that cellular states are shared across all GBM patients rather than specific driver mutations relevant only to specific patient subpopulations. This concept for targeting GBM is referred as ‘state selective lethality (SSL)’ (Nicholson and Fine, 2021).

In this study, we used single-cell RNA sequencing on glioma GSCs before, during, and after treatment with 13 diverse anti-glioma drugs to examine therapy-induced cell state changes. Most drug-induced state shifts reverted to baseline after drug removal, highlighting GSC plasticity. Panobinostat, a HDAC inhibitor, induced the most notable transition from AC-L to MES-L state. This epigenetically driven change, confirmed in barcoded GLICO tumors (Linkous et al., 2019), showed that astrocyte-like cells are especially sensitive to panobinostat. Moreover, manipulation of mesenchymal transcription factor FOSL1 altered panobinostat’s efficacy. Finally, combining drugs that drive opposite state transitions synergistically reduced GSC viability, suggesting a promising new multi cell-state targeted therapeutic approach.

## Results

### GBM cellular states are modulated by anti-tumor drugs

To explore whether therapeutic stress has an impact on cellular state distribution, glioma stem cells (GSCs) lines GSC-810 and GSC-320 growing as gliomaspheres were treated with the LD50 dose (Figure S1A) of 13 distinct anti-tumor agents. Drugs were selected based on their diverse mechanisms of action (MOA) and potential clinical relevance (Table 1). Single-cell RNA sequencing (scRNA-seq) was conducted before and after 72 hours of drug exposure and following a 14-day drug washout period to evaluate the cellular responses in treated and recovered cells (Figure 1A). Experiments were conducted across 5 experimental batches, with each batch containing an untreated baseline control – to account for the known plasticity of GSCs – and a DMSO vehicle control. Initially, cells were clustered and projected as a UMAP, and control cells (untreated baseline and DMSO conditions) were compared to uncover any vehicle-driven effects (Figure S1C). Based on this comparison, clusters 6 and 9—composed of approximately 90 percent DMSO 2-week-washout cells from GSCs 810 and 320, respectively—were removed, and newly filtered UMAPs were generated (Figure 1B). Inspection of projected UMAPs showed clear separation between the 810 and 320 genotypes, and highlighted panobinostat as the drug that caused the greatest transcriptional shift in both cell lines (Figure 1B). As expected, cell cycle phase signature scoring revealed decreases in the fraction of proliferating cells for the majority of drugs (Figure S1B). Curiously, we did not see a reduction in proliferation upon treatment with TMZ, presumably reflecting intrinsic GBM temozolomide resistance almost uniformly seen at the time of tumor recurrence (Liu et al., 2006). During the two week washout period, cells regained their proliferative capacity. We next performed principal component analysis on pseudobluk RNAseq samples to query how readily samples returned to their baseline transcriptional phenotypes after a two-week drug washout period (Figure 1C). For both GSC cell lines we observed separation of the drugged conditions across PC3, whereas vehicle-treated and washout conditions clustered nearer their baseline controls, re-emphasizing GSC’s remarkable plasticity.

**Figure 1:**
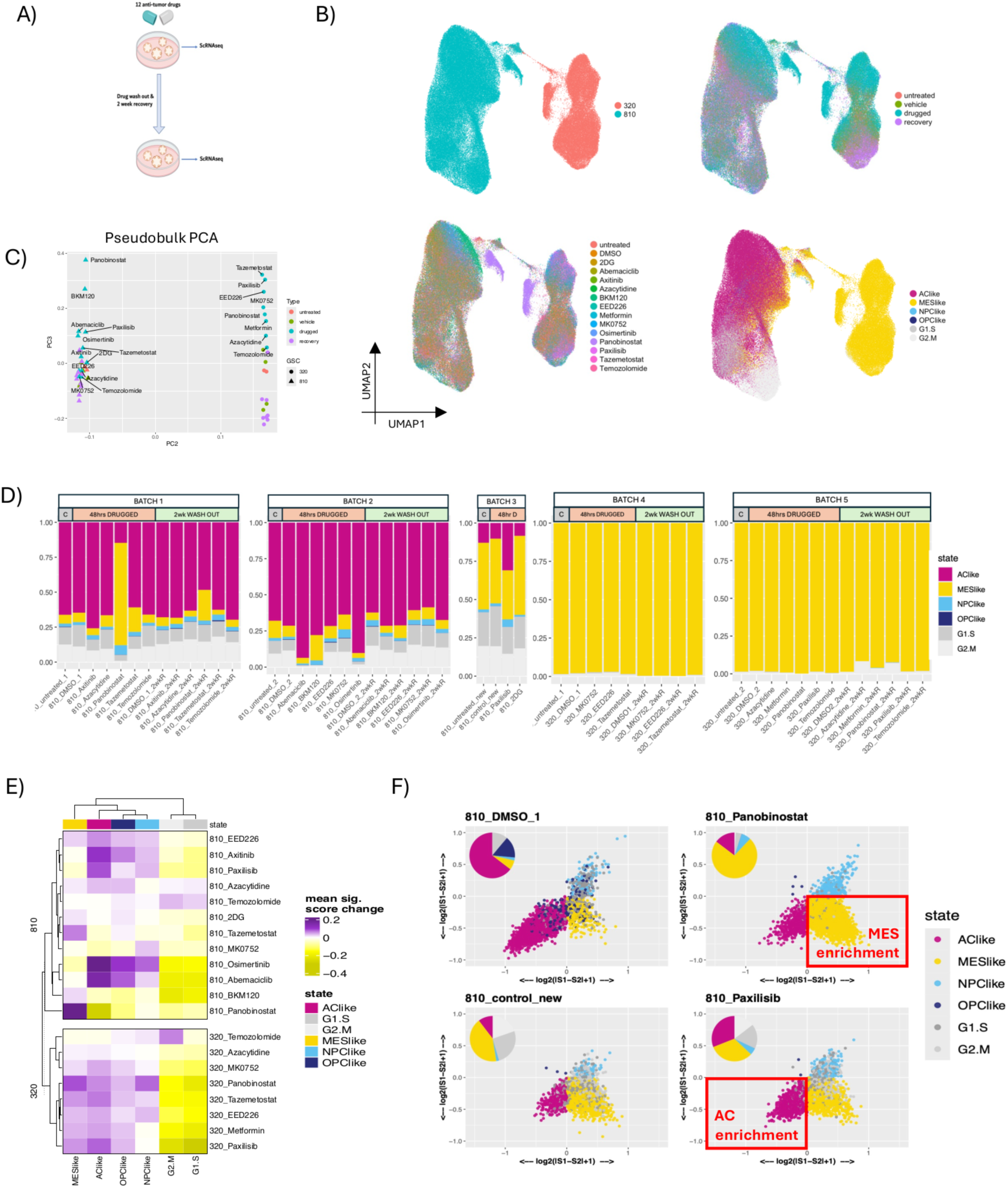
Anti-tumor drugs induce a significant, but reversible change in cell state distribution at single-cell level. a) Experimental schematic (left). b) UMAP projections of cells across all conditions colored by GSC genotype (top left), sample type (top right), drug condition (bottom left) and assigned Neftel cell state (bottom right) c) Principal component analysis of pseudobulk RNAseq samples d) Barplots displaying cell state composition, samples are seprated according to experimental batch, with each batch containing its own untreated and veichle (DMSO) controls. e) Heatmaps showing changes in mean Neftel state scores at 48hrs post-druggong relative to vehicle controls. f) 2D cell state quadrant plot (as defined in Neftel et al (ref)) of 810’s 72hrs post0treatment with panobinostat or paxilisib as well as their batch-specific vehicle controls. Quadrants: AC-L (bottom left), MES-L (bottom right), OPC-L (top left), and NPC-L (top right). Cells are colored by their assigned cell type inset Pie charts depict the percentages of the six cellular states in each group.

**Table 1:** Panel of 13 drugs used in this study.

| Drugs | IC50 Dose | Mechanism of Action | Cell state transitions (72 h treatment) | Cell state transitions (2 week recovery) |
| --- | --- | --- | --- | --- |
| Temozolomide | 100 uM | DNA alkylating agent | No alteration in cell state proportions | No alteration in cell state proportions |
| panobinostat | 50 nM | HDAC inhibitor | AC—> MES | Revert to baseline proportions |
| Tazemetostat | 50 uM | EZH2 inhibitor | MES enrichment | Revert to baseline proportions |
| Axitinib | 0.5 uM | VEGF receptor blocker | AC enrichment | Revert to baseline proportions |
| Azacytidine | 2.5 uM | DNMT inhibitor | No alteration in cell state proportions | MES enrichment |
| Osimertinib | 1 uM | EGFR inhibitor | MES —> AC | Revert to baseline proportions |
| Abemaciclib | 1 uM | CDK4 inhibitor | MES —> AC | Revert to baseline proportions |
| EED226 | 50 uM | PRC2 inhibitor | AC depletion | AC depletion |
| MK0752 | 50 uM | Gamma secretase inhibitor | AC—>NPC | Revert to baseline proportions |
| BKM120 | 25 uM | PI3K inhibitor | AC—> MES | Revert to baseline proportions |
| paxalisib | 500 nM | PI3K inhibitor | MES —> AC | N/A |
| 2DG | 1.5 mM | Glycolysis inhibitor | MES enrichment | N/A |
| Metformin | 6 mM | Antihyperglycemic agent | No alteration in cell state proportions | MES enrichment |

We next assigned GSCs to Neftel cell states (Neftel et al., 2019) based on signature scoring and evaluated the effect of drug exposure on GBM cellular states (Figure 1B, 1D). For GSC-810, which showed a balanced baseline distribution of GSC cell states, we identified three major groups of drugs; i) drugs that had little effect on cell state distribution, including Temozolomide, clinical standard of care for GBM, ii) drugs that induce a shift towards the MES-L state, most notably the histone deacetylase panobinostat, but also Tazemetostat to a lesser extent, and iii) drugs that polarized cells away from the MES-L state towards the AC-L state including paxalisib, Abemaciclib and Osimertinib. Notably, for all drugs that induced state changes, by the two-week washout time point, cells had relaxed towards the baseline distribution of their untreated control. In contrast to the varied baseline state distribution of GSC-810s, the GSC-320 cell line was very strongly polarized to the mesenchymal phenotype at baseline, and comparable state changes upon drugging were not evident (Figure 1D). However, GSC cell states are not distinct entities, but rather a set of continua of transcriptomic states defined by signature scoring and as such it is possible that a cell retains an overall MES-L assignment whilst its underlying signature scores changes. To test this theory, we computed mean signature scores changes relative to vehicle controls for all drug conditions (Figure 1E). Here, we observed matching state signature score changes in the 810s and 320 GSCs, with the panobinostat showing the greatest increase in the MES-L signature, and Paxilisisb showing the greatest increase in AC-L signature score. Since these two drugs also drove large state changes in the GSC-810s (Figure 1F), they were selected for further experimentation. Taken together, these findings highlight the dynamic nature of GBM cellular states and their capacity for plasticity in response to anti-tumor treatments.

### Panobinostat treatment show distinct cell state sensitivity through phenotype plasticity *ex-vivo*

As often occurs in many epithelial cancer types, shifts in the genotypes of drug-resistant cell populations are a result of drug-induced genetic clonal selection. Although clinical and genetic data from patients with recurrent GBM do not support predominant genetic clonal selection at the time of patient relapse, we wanted to rule out the possibility that this was occurring in our in vitro model system. To do so, we evaluated copy number alteration in treated GSCs and found no evidence of genetic clonal selection, suggesting that phenotypic plasticity accounts for the observed changes in cellular states (Figure S2A-B). To verify this and to investigate cellular state plasticity at single-cell resolution under therapeutic stress, we utilized a genetic bar-coding clonal tracking (Guo et al., 2019) approach in our *ex-vivo* GLICO model.

Our prior data have shown that the GLICO microenvironment provides GSCs with enhanced access to diverse cellular states compared to traditional in vitro/2D cultures, potentially contributing to the increased drug resistance observed in GLICO (Pine et al, 2020). To test whether drug-mediated state transitions are retained in GLICO, and in an independent cell line, 1206-GSCs, we focused on panobinostat, the drug that caused the most significant impact on cell state changes and plasticity in our drug screening panel (Figure 1C-D). We repeated the drug treatment in GLICO but using CellTaged (Guo et al., 2019) GSCs to concurrently track clonal history and cell state identity (Figure 2A-B). We identified a variety of clones, with clones 1, 4, 6, 8, and 13 accounting for half of the total population under both conditions (Figure 2C). The distribution of cell states following exposure to panobinostat revealed an increase in MES-L cells at the expense of AC-L cells, mirroring observations from 2D cultures (Figure 1D). Notably, clonal dynamic analysis before and after treatment revealed specific trends: a substantial number of AC-L clones transitioned to MES-L clones after treatment (Figure 2D). Additionally, several AC-L dominant clones (Clones 4, 8, and 13) were reduced in cell number following, whereas others, such as Clone 1 and Clone 6, expanded in size, predominantly through the proliferation of MES-L and NPC-L cells, respectively (Figure 2E). In order to rule out panobinostat-mediated genetic clonal selection rather than single cell cellular state plasticity, we performed Whole Genome Sequencing (WGS) following the same experimental scheme, which showed no evidence of large scale genetic clonal selection through bulk copy number alteration analysis (Figure S2A). To test whether more subtle changes in copy number heterogeneity could explain dynamic changes in barcode composition, we performed copy number analysis on scRNAseq data. Results showed evidence of genetic clonal heterogeneity with three genetic subclones detected (Figure S2B). However relative proportions of genetic clones were largely unchanged after panobinostat treatment (Figure S2C-D). Comparable analyses of our in vitro screening data revealed consistent results without systematic changes in genetic subclone composition post-drug treatment in either 810 or 320 GSCs (Figure S2E-F). Therefore, the observed clonal dynamics can be explained mostly at the phenotype level. Overall, these shifts reflect the transcriptomically mediated cell state plasticity of GSCs, explaining the observed transition from AC-L to MES-L cell states, and highlighting the increased vulnerability of AC-L cells to panobinostat (Figure 2F).

**Figure 2:**
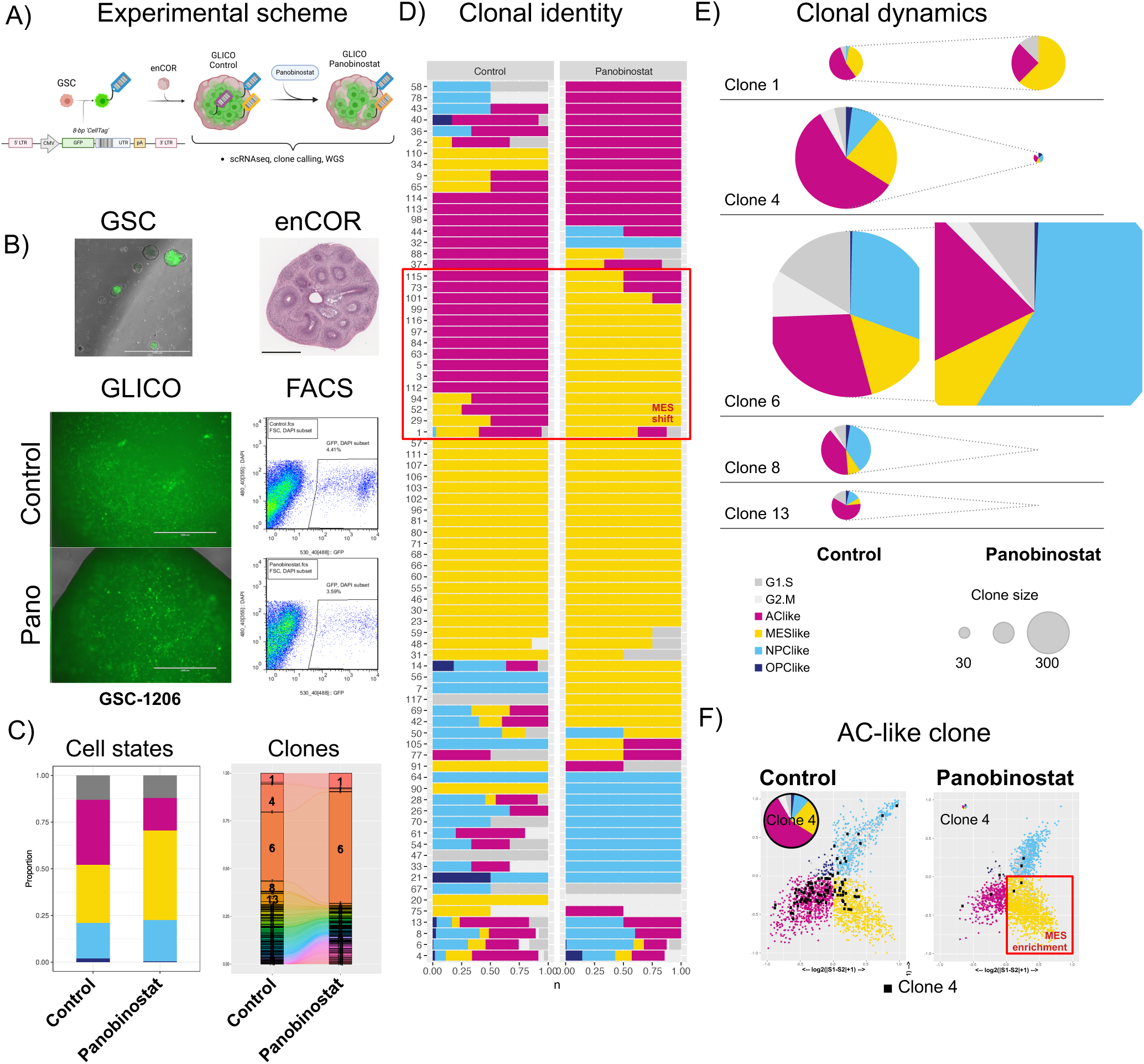
Barcoded cells show differentiated cell state sensitivity through phenotype plasticity in GLICO. a) Experimental scheme for barcoded GSCs in GLICO treated with panobinostat. b) Representative pictures of GSCs, enCOR, and GLICOs treated or not with panobinostat and sorting for GSCs GFP+ DAPI-. c) Clone calling bar plots of barcoded cells before and after treatment, where each color represents a single clone (left) and Cellular State distribution between conditions (right). d) Barplots displaying cell state composition for each clone before and after panobinostat treatment. e) Clonal dynamics plots of individual clones before and after treatment for the five major identified clones in (C). Pie charts show cell state proportion on each clone and pie size represents clone size. f) Cell state quadrant plot. Quadrants: AC-L (bottom left), MES-L (bottom right), OPC-L (top left), and NPC-L (top right). Cells are colored by their assigned cell type, with AC-L clone 4 highlighted in black. Inset Pie charts depict the percentages of the six cellular states in each group for Clone 4 and pie size represents clone size before and after treatment.

### FOSL1-mediated cell-state redistribution to MES-L state affects sensitivity to Panobinostat

To more directly evaluate the effects of cellular states on therapeutic stress, we investigated whether genetic perturbation of a transcription factor known to mediate the shift to the MES-L state could alter treatment responsiveness. Specifically, we manipulated FOSL1 by either increasing or decreasing its expression prior to panobinostat treatment (Figure 3A). Consistent with our hypothesis that the transition to the MES-L state represents a mechanism of panobinostat resistance, cells overexpressing FOSL1,which promotes the MES-L state transition (Chen et al., 2022) (Figure 3C), exhibited significantly increased resistance to the cytotoxic effects of panobinostat. In contrast, cells with reduced FOSL1 expression, resulting in fewer MES-L cells (Marques et al., 2021) (Figure 3C), showed increased sensitivity to panobinostat compared to uninduced cells (Figure 3D). Correspondingly, apoptosis levels post-panobinostat treatment varied with FOSL1 expression: higher FOSL1 levels protected against apoptosis, while FOSL1 knockdown (KD) enhanced apoptotic responses (Figure 3E). Overall, these findings validate that manipulating cell state distribution, specifically through the modulation of FOSL1, significantly influences sensitivity to panobinostat, underscoring the effectiveness of targeted cell-state distributions in enhancing treatment responsiveness.

**Figure 3:**
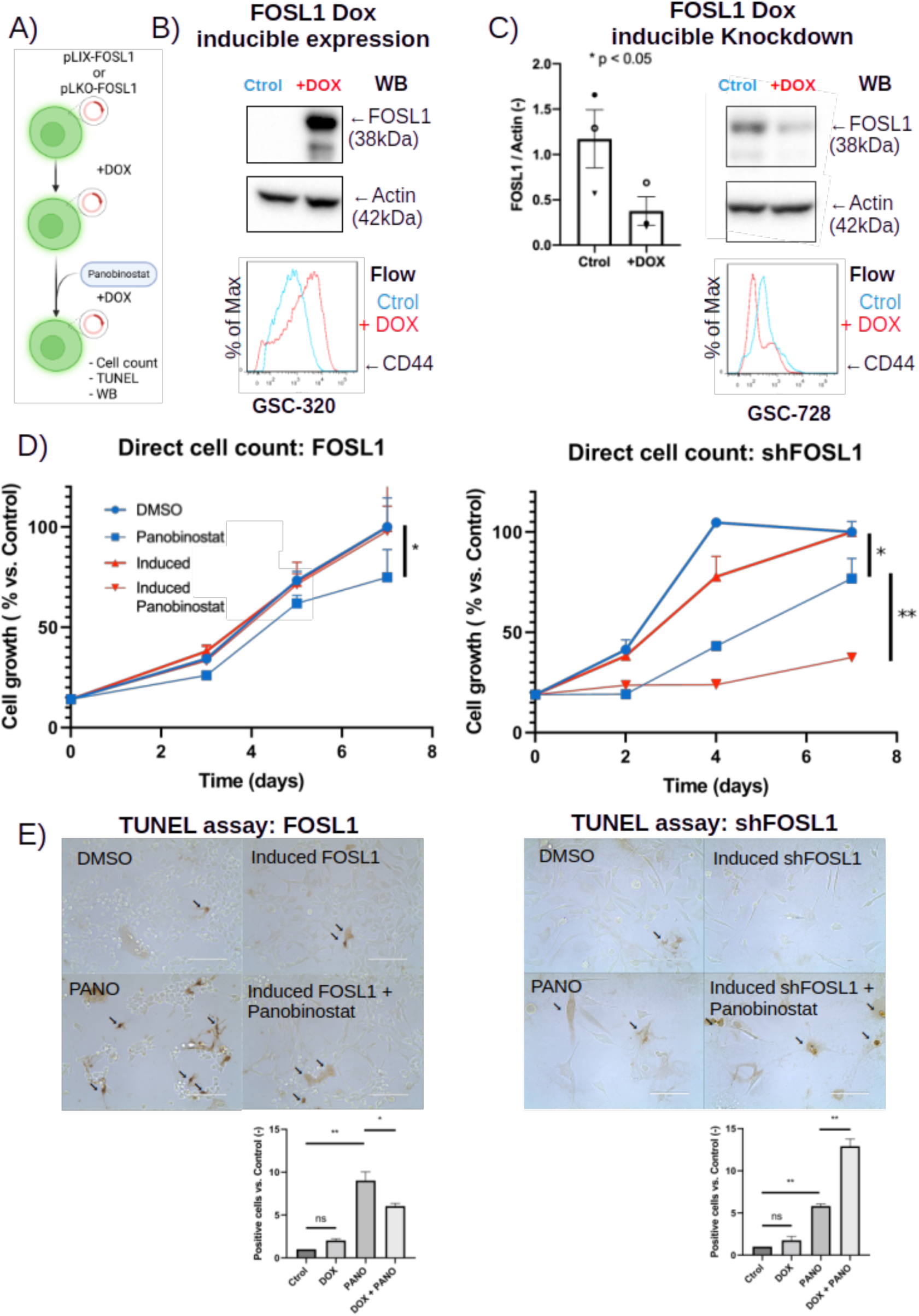
Genetic perturbations trap cells into specific sensitive states. a) Schematic overview of GSC-320 inducible over-expression (OE) and GSC-728 inducible knock-down (KD) of FOSL1, followed by panobinostat treament. B) FOSL1 expression levels measured by Western blot. Actin expression level was used as protein loading control (top). Histograms of CD44 (MES-L state marker) expression measured by flow cytometry (bottom). GSC-320 cells transduced with doxycycline-inducible FOSL1 construct; DOX, GSCs treated with 2.5 μg/mL of doxycycline during 72 hours. C) FOSL1 expression levels measured by Western blot. Actin expression level was used as protein loading control (top). Histograms of CD44 (MES-L state marker) expression measured by flow cytometry (bottom). GSC-728 cells transduced with inducible KD FOSL1 construct DOX, GSCs treated with 2.5 μg/mL of doxycycline during 72 hours. D) GSCs growth curves for inducible FOSL1 OE (left) and (KD) cells treated or not with panobinostat. Viable cells were quantified by direct cell count using trypan blue exclusion. E) Transferase dUTP Nick End Labeling (TUNEL) assay for inducible FOSL1 OE (left) and KD cells (right) treated or not with panobinostat. Proportion of positive apoptotic cells, indicated by arrows, were counted per field and normalized to Control.

### Drug combinations targeting different cellular states demonstrate synergistic effects

We have previously postulated that cellular state plasticity might be a mechanism partially responsible for intrinsic glioma therapeutic drug and radiation resistance, a phenomenon that could be therapeutically manipulated using a strategy we called, “State Selective Lethality (SSL)” (Nicholson and Fine, 2021). In brief, SSL involves pushing cells with one drug into a cellular state more vulnerable to cytotoxic attack by a different drug. To explore this potential, we investigated whether combining panobinostat, a MES-inducing drug, with drugs that suppress the MES-L state could result in therapeutic synergy. GSCs were treated with panobinostat, paxalisib, and their combination, using a constant proportion ratio scheme. Drug response curves indicate that the combination treatment exhibited a greater effect than the individual treatments (Figure 4A). Interestingly, the improved efficacy is still observed after inducible expression of FOSL1 (Figure 4B). Combination Index (CI) scores indicated that the combined treatment elicited a synergistic response compared to either drug alone, with improved synergism following FOSL1 induction, as denoted by lower CI values (Figure 4C, S3A). Similarly, the combination of panobinostat with other MES-suppressing agents, such as Abemaciclib and Osimertinib, resulted in a more pronounced reduction in cell viability compared to the effects of the individual drugs (Figure 4D, S3B). Next, to further validate the principle of state-selective lethality, we compared these findings with results from two drugs that both drive cells toward the MES-L state - panobinostat and 2DG. In this scenario, the combination therapy did not exhibit any advantage over individual treatments (Figure S3C). Moreover, inducible expression of FOSL1 significantly diminished the inhibitory effect of 2DG by approximately half, while others MES-inducers compounds showed either a recovery or no increase benefit over inducible FOSL1 expression (Figure S3C).

**Figure 4:**
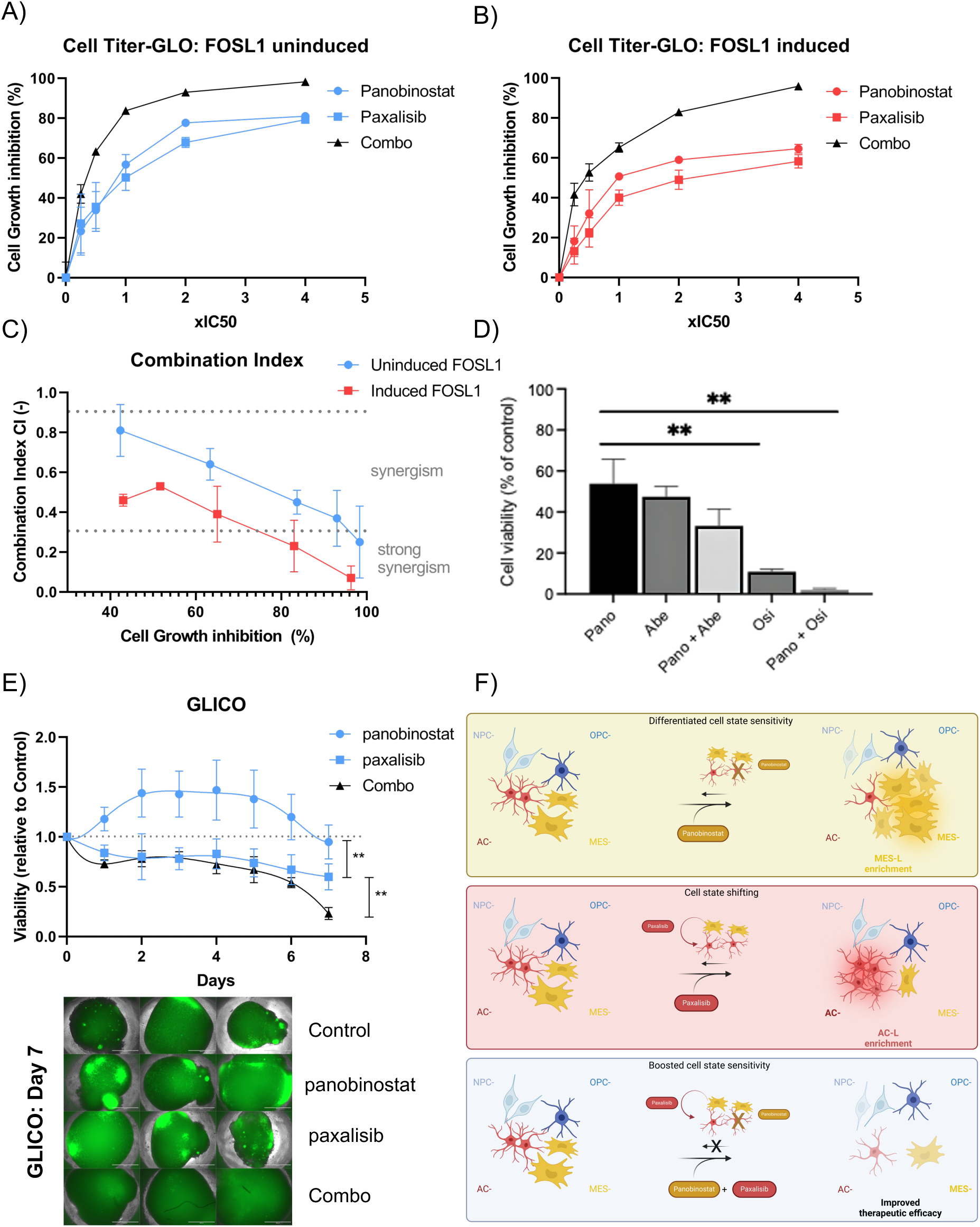
Combination treatment of AC-L and MES-L inducers produce a synergistic effect. a) Cell Titer Glo experiments for panobinostat, paxalisib and combination (Combo) treatments on uninduced 810 FOSL1 cells. Each curve point represents the effect on Cell Growth inhibition of a fraction of the IC50 for a given treatment. b) Cell Titer Glo experiments for panobinostat, paxalisib and combination (Combo) treatments on induced 810 FOSL1 cells. Each curve point represents the effect on Cell Growth inhibition of a fraction of the IC50 for a given treatment. c) Combination index plots vs Cell Growth Inhibition curves for uninduced and induced 810 FOSL1 cells, where CI = 1 indicates agonism, CI > 1 antagonism and CI < 1 synergism. D) Cell Titer Glo experiments for panobinostat, Abemaciclib, Osimertinib and their combination on uninduced 810 FOSL1 cells. E) 810-NlucP-GFP GLICOs treated with the Cmax of panobinostat, paxalisib and combination (Combo) over the period of 7 days. Relative luminiscence fold-change over time is normalized to Control (DMSO). ** p < 0.01 two-way ANOVA (top). Representative images of treated GLICOs at Day 7. Bar: 1000um (bottom). F) Graphical abstract.

Finally, we assessed the proposed state-selective lethality concept in our ex vivo model using a sensitive luminescence-based assay to track cellular growth longitudinally. GLICOs were treated with panobinostat, paxalisib, or their combination. Interestingly, panobinostat alone did not significantly reduce cell growth compared to control, whereas paxalisib exhibited notable inhibitory effects. Importantly, the combination treatment showed statistically significant inhibition compared to either single agent from day 4 onward, with a four-arm longitudinal analysis (Mao, B. et. al, 2023) confirming synergy between panobinostat and paxalisib (Figure 4E, S3E).

These results underscore the potential efficacy of a SSL-like strategy in targeting different cellular states with combinatorial drug therapies to achieve superior therapeutic outcomes in GBM (Figure 4F).

## Discussion

Our study underscores the complex dynamics of GBM cellular state responses to anti-tumor treatments. *In vitro* drug screening of GSCs with 13 anti-tumor agents revealed varied changes in cell state distributions, which predominantly shifted the balance between MES-L and AC-L states. After a two-week recovery post-drug washout, cells relaxed to their baseline cell state distributions, suggesting that drug-induced shifts towards a ‘refuge’ state are temporary. We have previously postulated that rational design of therapies that counteract adaptive cell state transitions may reduce drug resistance, a concept we refer to as ‘state selective lethality (SSL)’ (Nicholson and Fine, 2021). Here, we provide proof of concept for SSL by demonstrating that exogenous manipulation of cellular states through FOSL1, a key mediator of the MES-L state (Chen et al., 2022), can modulate the resistance to panobinostat by limiting or increasing access to the MES-L ‘refuge’ state. Further, we showed that combinatorial treatment of GSCs with panobinostat and drugs inducing AC-L refuge state (paxalicib, abemaciclib or osimertinib) elicited synergistic responses.

The dynamic transitions of cellular state response to drugs could be categorized intro three groups, those inducing a MES-L shift (panobinostat, 2DG, BKM120 and Tazemetostat), those inducing a AC-L shift (paxalisib, Abemaciclib and Osimertinib) and those that did not shift cellular states. TMZ’s membership in this final group is intriguing, and it is tempting to speculate that the inability of GSCs to escape Temozolomide by cell state transitions contributes to its clinical efficacy. An unexpected finding was the differential effects of two PI3K inhibitors tested: BKM120 and paxalisib. BKM120 caused a shift towards the MES-L state, while paxalisib induced an AC-L shift. This discrepancy may be attributed to their differing efficacies in targeting PI3K subunits and/or to the fact that paxalisib is also a potent mTOR inhibitor. Further work will be required to evaluate whether other drugs with similar mechanisms of action cause comparable cell state shifts, possibly aided by high-throughput screens using fluorescent reporters of GBM cellular states.

Drug-induced changes in GSC cell state proportions could be explained by variable cell state sensitivities and rates of cell death; plastic state transitions from sensitive to refuge states, or some combination of the two. Our CellTag barcoding experiments suggests that both mechanisms contribute to the observed changes in cell state distributions as we saw depletion of AC-dominated clones and increases in MES dominated clones as well as plastic transitions of cell states within clones. The balance between the variable rate of cell death and plasticity may govern the balance between cytostatic or tumor-reducing treatments and warrants further investigation. Whilst we cannot completely exclude the possibility that genetic, rather than epigenetic state selection drives the transcriptional changes we see upon drug treatments, our copy number and bar coding analysis suggested that the contribution of genetic clonal selection is limited. This is consistent with large-scale longitudinal studies in GBM, which have found little evidence of clonal evolution upon recurrence (Barthel et al., 2019).

One of the theoretical advantages of SSL therapeutic approach is its focus on cellular states shared across all GBM patients an clonal populations, rather than on specific driver mutations and the highly variable clonal genetic landscapes relevant only to a subset of intratumoral clones and a specific subpopulation of patients (Suva and Tirosh, 2020).In this regard, it was encouraging that the drug synergies predicted by our analyses of 810 and 1206 GSCs also proved effective on 320 GSCs, which were dominated by the mesenchymal phenotype. Similarly, by treating FOSL1-induced 810 GSCs with panobinostat and paxalisib we showed that drug synergy was maintained in a situation in which a previously sensitive cell line underwent a mesenchymal shift. This mimics the natural transcriptional progression of many GBMs, and suggests our approach may be applicable to recurrent GBMs.

We have previously shown that culturing GSCs in 2D limits their access to more stem-like NPC/OPC phenotypes compared to models with more complex microenvironments such GLICO co-cultures or orthotopic xenografts (Pine et al., 2020). It was notable that none of the drugs we tested in vitro induced a transition to either stem-like state. While we did recapitulate the in vitro–observed synergism between Panobinostat and Paxalisib using GLICOs, it is possible that the potential for identifying SSL drug-synergistic combinations extends beyond those predicted by our initial 2D screen. Other studies demonstrating synergy between drugs targeting stem-like and mesenchymal cell populations support this hypothesis (Garnier et al., 2019; Wang et al., 2021; Lu et al., 2025). Thus, future studies will need to implement large-scale single-agent and combination testing in models such as GLICO, which allow access to a broader representation of the clinically observed cellular states than is captured in the in vitro screen.

Taken together, our study highlights the remarkable plasticity and variability of GSC cellular state responses to anti-tumor therapies. Through genetic perturbation of cellular states and rational design of combinatorial treatments of drugs that induce opposing cell state transitions, we provide proof-of-concept data that state-selective lethality is a potentially promising strategy for overcoming treatment resistance and achieving more effective GBM therapies. Future research should continue to explore these mechanisms and their potential clinical applications.

## Acknowledgements

We would like to thank the patients whose sample donations were used in this study. We also thank Alicia Alonso and Yushan Li from the WCM Epigenomics Core, Sharmaine Griffith-Baker from LBPMC Core, the WCM Genomics Core and the WCM Flow Cytometry Core for their support. Work in H.A.F.’s laboratory was supported by an NIH Director Pioneer Award (1DP1CA228040-01).

## Author Contributions

S.M.C., R.S., J.G.N. and H.A.F. designed the study, interpreted the results, and wrote the manuscript. S.M.C., R.S. performed experimental work. S.M.C. and J.G.N. performed computational analysis. YW, CH, AM and SG contributed to organoid work and provided experimental and technical support. H.A.F. supervised the work.

---

**Supplementary Figure 1.**
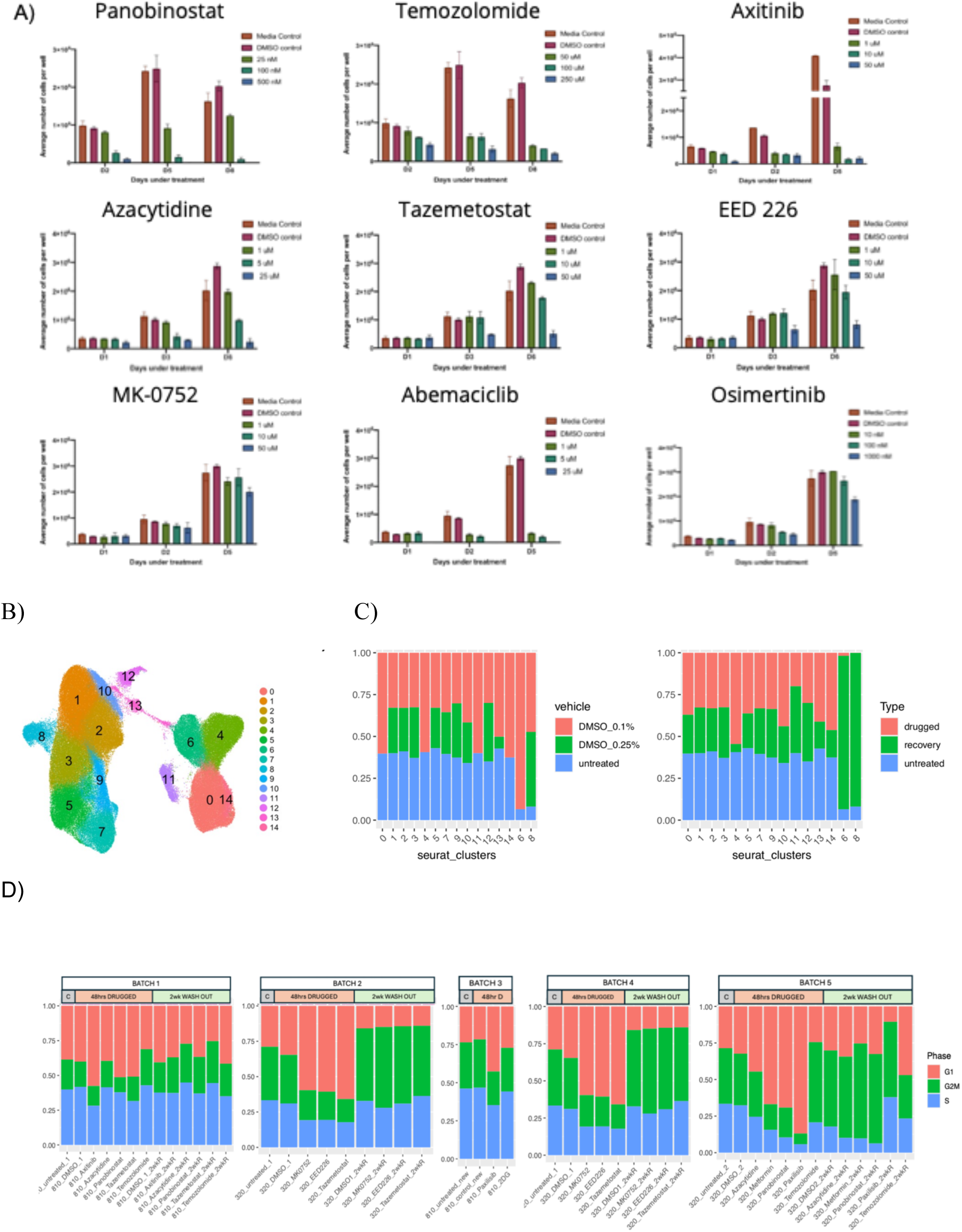
a) Dose response assays of nine drugs in 320 GSCs to determine IC50 dose for scRNAseq experiments (IC50 of other drugs taken from literature or calculated elsewhere) b) UMAP projection of all cells, colored by cell cluster c) Barplots showing cluster composition of control cells (untreated and vehicle 72hrs and 2week washout), left shows vehicle vs control conditions and right shows samples type. d) Barplots displaying cell phase composition, samples are separated according to experimental batch, with each batch containing its own untreated and veichle (DMSO) controls.

**Supplementary Figure 2:**
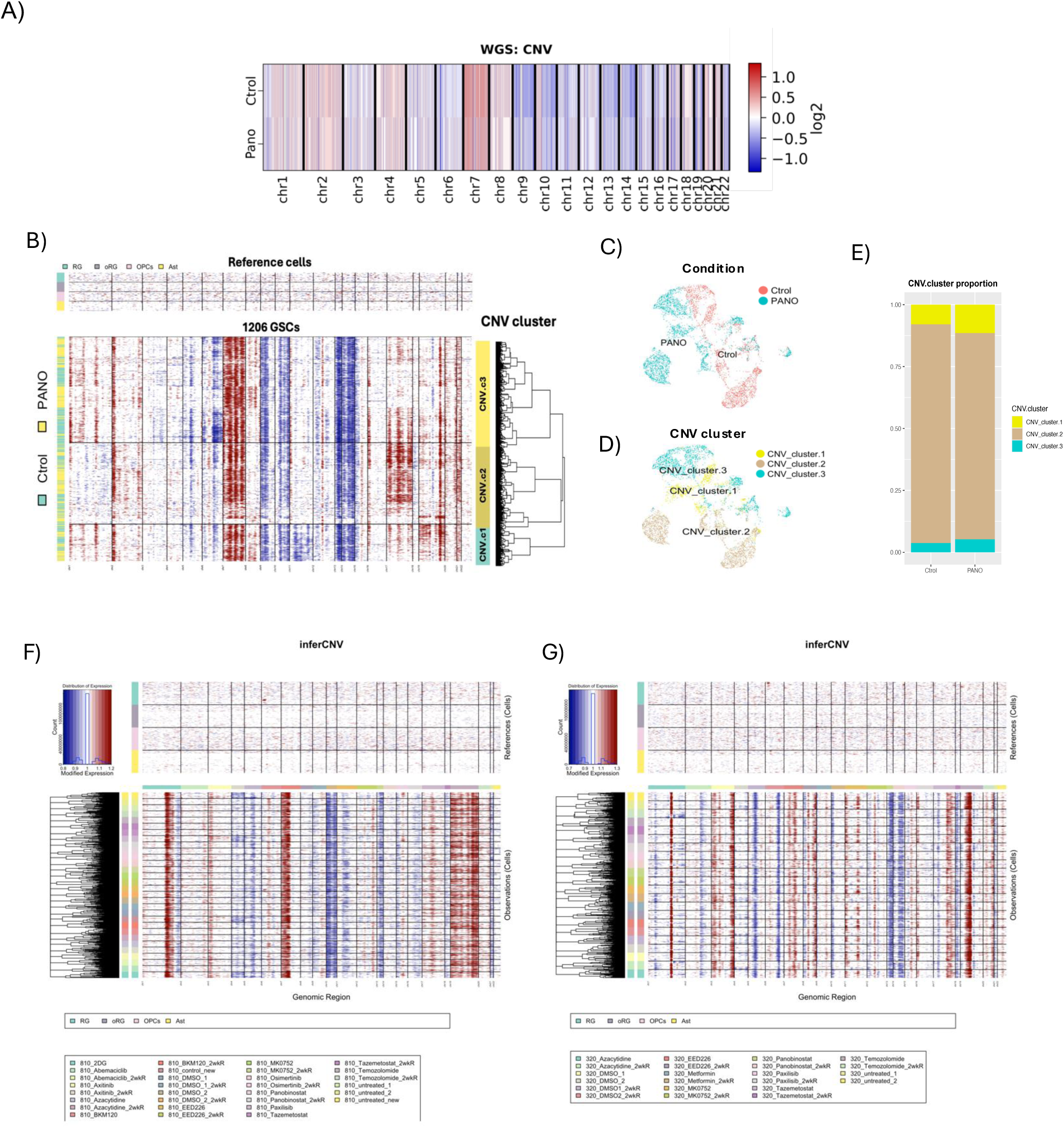
a) Copy number variation comparison from WGS of GSC-1206CellTag line in GLICO treated or not with panobinostat. b) inferCNV comparison from scRNAseq of GSC-1206CellTag line in GLICO treated or not with panobinostat. c) inferCNV analysis for all drugs in GSC-320. d) inferCNV analysis for all drugs in GSC-810.

**Supplementary Figure 3:**
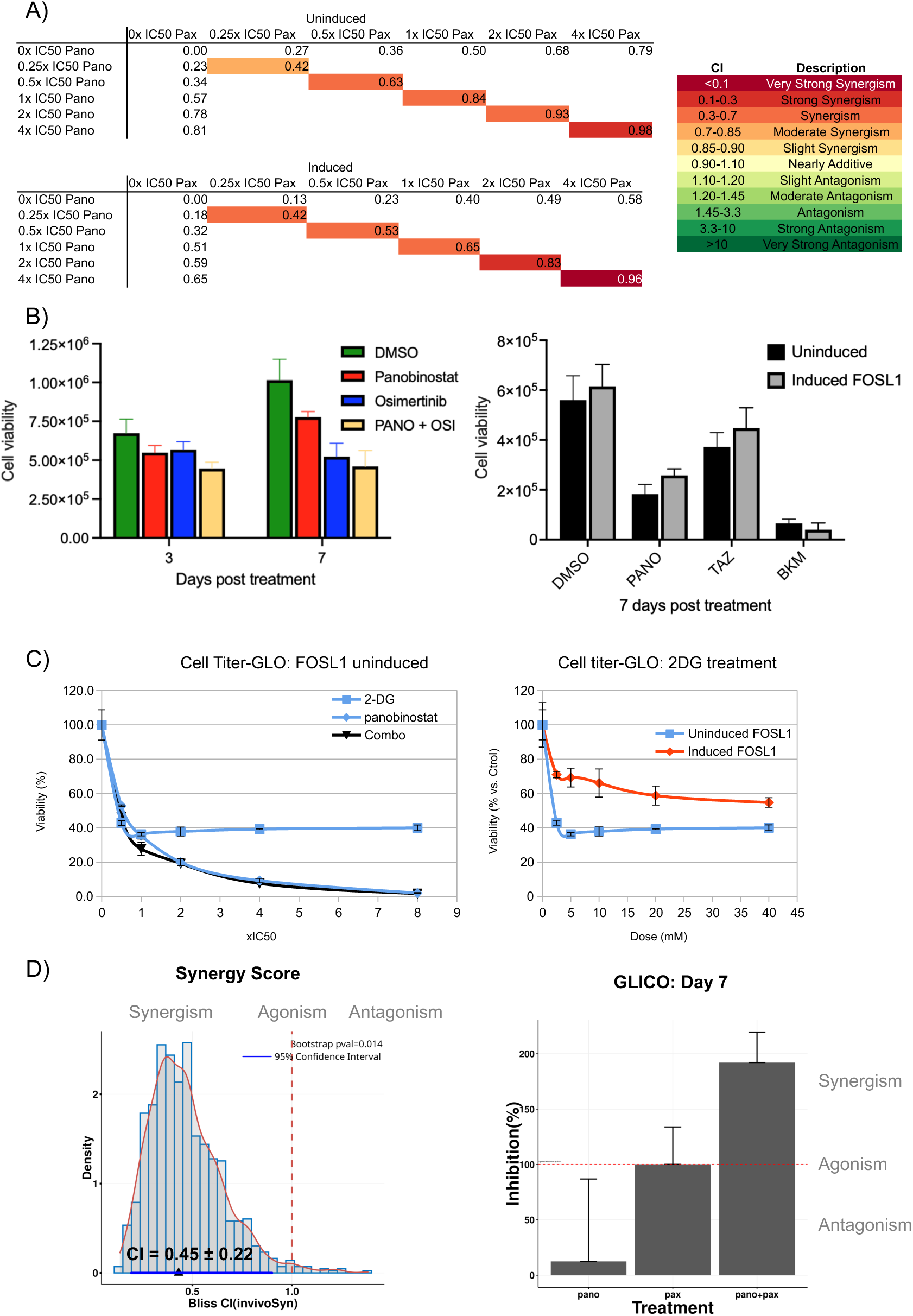
A) Tables with Fraction affected (Fa) values of diagonal combination treatments of panobinostat and paxalisib on a constant proportion scheme for Uninduced (top) and Induced FOSL1 (bottom). Colors in the diagonal represent the synergistic effect of the CI values calculated for each Fa. B) panobinostat, Osimertinib and their combination on GSC-320 uninduced FOSL1 cells (left). Cell Titer Glo experiments for panobinostat, tazemetostat and BKM120 on induced 810 FOSL1 cells (right). C) Cell Titer Glo experiments for panobinostat, 2DG and combination (Combo) treatments on uninduced 810 FOSL1 cells. Each curve point represents the effect on Cell Growth inhibition of a fraction of the IC50 for a given treatment (left). Cell Titer Glo experiments for 2DG treatment on uninduced and induced 810 FOSL1 cells (right). E) Bliss Combination Index (CI) of panobinostat and paxalisib calculated for the whole GLICO experiment (left) and Day 7 (right). (CI = 1 → additive, CI > 1 → antagonism, CI < 1 → synergy).

## STAR★Methods

### RESOURCE AVAILABILITY

#### Lead Contact and Materials Availability

Further information and requests for resources and reagents should be directed to and will be fulfilled by Howard A. Fine.

#### Data and Code Availability

Single-cell data have been deposited at GEO and are publicly available as of the date of publication. Accession numbers are listed in the key resources table.
This paper does report original code.
Any additional information required to reanalyze the data reported in this paper is available from the lead contact upon request.

### EXPERIMENTAL MODEL AND SUBJECT DETAILS

#### Patient-Derived GSCs

Following informed consent, tumor samples classified as glioblastoma, based on the World Health Organization (WHO) criteria, were obtained from patients undergoing surgical treatment at the National Institutes of Health (NIH) or from Weill Cornell Medicine/New York Presbyterian Hospital in accordance with the appropriate Institutional Review Boards. Within 1-3 hours after surgical removal, tumors were washed in PBS and enzymatically dissociated into single cells. Tumor cells were cultured in NBE medium consisting of neurobasal medium (Thermo Fisher Scientific), N2 and B27 supplements (Thermo Fisher Scientific), and human recombinant bFGF and EGF (25 ng/mL each; R&D Systems) plus Heparin sodium and L-Glutamine. Regular mycoplasma screening was performed using the MycoAlert Detection Kit (Lonza Inc.).

#### Human ESCs

NIH-registered human H1 (WA01) or H9 (WA09) embryonic stem cells were purchased from WiCell Research Institute, Inc. and maintained in mTeSR1 medium (STEMCELL Technologies).

### METHOD DETAILS

#### Glioma Cerebral Organoid (GLICO) Generation

Overview of the experiment is provided in Figure 1A. Cerebral organoids and co-culture with GSCs were generated as described in (Linkous et al, 2019). Briefly, GSCs were transduced with pLenti PGK GFP Blast (w510-5) (Addgene) and were cultured under blasticidin selection in NBE medium. Organoids were plated one per well in a 96-well round bottom plate, excess medium was removed, and 50,000 GSCs that stably express GFP in 150ul of NBE were added to each well. After stationary incubation at 37°C for 24hr, GLICOs were washed in PBS and transferred to a new 6-well plate with 4ml of organoid differentiation medium. Organoids were maintained on an orbital shaker for up to 10 days at 37°C. Representative images were acquired using an Evos FL Auto imaging system (Carlsbad).

#### Single-cell dissociation and sorting

GLICOs were dissociated into single-cell suspensions using the Papain Dissociation System following manufacturer’s instructions (Worthington Biochemical, LK003150). 2D cultures were dissociated following standard passaging protocols using Tryple. Cells were filtered through a 40um strainer and resuspended on ice-cold PBS/2%BSA. GFP-positive/DAPI–negative cells were sorted by FACS (Becton-Dickinson Aria II) and collected on PBS/0.04%BSA for single-cell, RNA- or Whole Genome-Sequencing. Cell viability was checked on every step of the process by Trypan Blue Exclusion test.

#### scRNA-sequencing processing

scRNA-seq libraries were prepared with the Chromium Single Cell 3′ Library & Gel Bead Kit v.2 (10×Genomics, PN-120237), and sequenced on a NextSeq 500 instrument (Illumina). The Cell Ranger 2.0.1 pipeline was used to align reads to the GRCh38 human reference genome and produce count matrices for downstream preprocessing and analysis using the Seurat v5.0 R package[3]. For quality control, cells with fewer than 1000 genes detected, fewer than 2250 reads, or greater than 25% mitochondrial gene expression were removed. Genes detected in fewer than 3 cells were excluded from analyses. Expression values were library size corrected to 10,000 reads per cell and log1p transformed, with Principal component analysis (PCA) performed on the scaled data for the top 2000 variable genes. Batch correction was performed on principal components using Harmony [4]. Uniform Manifold Approximation Projection embeddings, Nearest Neighbors and cell clusters were then calculated in harmony-corrected PCA space using the default settings of Seurat’s RunUMAP(), FindNeighbors(), and FindClusters() functions. Initial cluster composition was compared to uncover any vehicle driven effects (Figure S1 A-B), with clusters 6 and 9, respectively made up of >90% DMSO 2week-washout cells from GSCs 810 and 320, removed before re-clustering cells as before with default function settings. For reference signature scoring, average gene module expression was calculated for each single cell, subtracted by the aggregated expression of a random control set of features selected from the same average expression bins as the query genes[5]. Inbuilt cell cycle phase signatures were taken from Seurat v5.0 and GSC cell state signatures were taken from and visualized as in Neftel et al., [6]. For mean cell state signature scoring, the difference between sample averages and their batch- and DMSO concentration-matched concentration were reported. For copy number analysis of barcoded 1206 GSCs inferCNV [7] was used in tumor sub-clustering mode with a lieiden resolution of 0.0001. CNV cluster annotations were transposed to the Seurat object and cluster propotion changes in barcoded clones of at least 5cells were visualized. For copy number analysis of in vitro data, a random subset of 1000 cells from each sample was analysed with inferCNV using default settings.

#### Whole Genome sequencing (WGS)

The WGS data used in this study were generated using PacBio Revio sequencing for low input samples following manufacturer’s protocol (Wang et al., 2023). Briefly, high molecular weight (HMW) DNA from samples was extracted and quantified to meet the minimum input requirement of 500 ng. SMRTbell libraries were prepared using the PacBio SMRTbell Express Template Prep Kit 2.0, ensuring uniform coverage and minimal DNA loss during library preparation. Libraries were sequenced on the PacBio Revio system to generate high-fidelity (HiFi) long reads, employing circular consensus sequencing (CCS) technology to achieve high accuracy across individual reads. Post-sequencing, data were processed through the PacBio Revio analysis pipeline, which includes base calling, CCS generation, and alignment to the reference genome using PBAlign.

#### Cell Viability and Apoptosis

GSCs were under drug treatment for the indicated time frame in the results section. Viable cell number were measured either by direct cell count using trypan blue exclusion or Cell-Titer Glo 3D (Promega), following manufacturer’s instructions. In GLICOs, GSC viability was inferred by the presence of stable luminescence expression (NanoLucPest, Promega). For apoptosis analysis, GSCs attached to matrigel coated Lab-Teks were treated with drug during 72h prior TUNEL assay kit, following manufacturer’s instructions (Thermo). The number of apoptotic cells were counted per field and normalized to the Control condition.

#### FOSL1 Inducible Expression

FOSL1 ORF was sub-cloned into an Inducible lentiviral expression backbone (Addgene #41395) using the Gateway technology. Product sequence was confirmed by Sanger sequencing. GSCs were transduced with the obtained plasmid and were cultured under Puromycin selection in NBE medium. For inducible FOSL1 expression, GSCs growing as 2D were treated with 2.5ug/ml of Doxycycline for 72 hours prior to further analyses or drug treatment, where Doxycycline induction was present during the whole experiment.

#### shFOSL1 Inducible Expression

shRNA oligos against FOSL1 ORF were sub-cloned into an Inducible lentiviral expression backbone (Addgene #21915). Product sequence was confirmed by Sanger sequencing. GSCs were transduced with the obtained plasmid and were cultured under Puromycin selection in NBE medium. For inducible FOSL1 expression, GSCs growing as 2D were treated with 2.5ug/ml of Doxycycline for 72 hours prior to further analyses or drug treatment, where Doxycycline induction was present during the whole experiment.

#### Western blot analysis

GSCs growing as neurospheres were washed twice with ice-cold PBS and then lysed with RIPA buffer (Thermo Scientific) with protease inhibitors. Samples were run in Novex™ NuPAGE™ 4-12% Bis-Tris SDS-PAGE (Invitrogen™), and the gels blotted to PDVF membranes (Invitrogen). Membranes were blocked for 1 h in 5% skim milk in PBS, 0.1% Tween-20. Membranes were then incubated with the first antibody overnight at 4 °C, and with a secondary antibody coupled to horseradish peroxidase [1 h at room temperature (RT)]. Detection was performed by chemiluminescence. Bands were digitalized with ChemiDoc MP Imaging System (Bio-Rad) and signal intensity was quantified with ImageJ software.

#### Flow cytometry analysis

GSCs were stained with CD44 antibody coupled to APC (#130–113–338, Miltenyi Biotec) followed the manufacturer’s cell surface flow cytometry staining protocol. Flow cytometry was performed using the BD Aria II Analyzer. Cell debris and dead cells were excluded from the analysis based on scatter signals and DAPI fluorescence.

### QUANTIFICATION AND STATISTICAL ANALYSIS

#### Cell state identification

To identify the cell state in a scATAC-seq cell, we first found the nearest scRNA-seq cell in joint CCA embedding using the Seurat FindTransferAnchors() method and assigned the matched cell’s transcriptome to the ATAC cell. The accessibility around a gene locus, quantified by a ArchR’s gene score, and the corresponding gene expression values were used as integration features. GBM cellular state was assigned by scoring the expression of gene markers, as defined in (Neftel et al., 2019), of the matched transcriptome for scATAC-seq cells, or the measured transcriptome for multiome cells.

#### Cellular Barcoding and Clonal Dynamics

Glioma stem cells (GSCs) were labeled using the CellTag library, following the protocol published by Kong et al. (2020). Briefly, a barcode whitelist was generated, and library complexity was verified through amplicon deep sequencing. GSC-1206 cells were transduced with the CellTagV1 lentivirus at a multiplicity of infection (MOI) of 3:1. Transduced cells were maintained for 10 passages to allow sufficient clonal expansion before initiating experiments. Clone calling and downstream analyses were performed using the CellTagR package.

#### Drugs interaction analysis

To determine the combination index (CI) of panobinostat and paxalisib, cells were treated with both drugs following a constant-ratio analysis. The combination ratio used was equal to the IC50 ratio for each drug. Two serial dilutions below and above the IC50 value were used for each mixture. Effects of all the mixture points were displayed using the form CI vs. Fa, where Fa is fraction affected and represents the respective proliferation inhibition parameters (e.g., a Fa of 0.5 is a proliferation inhibition of 50%). CI plot values were obtained from three different experiments using Compusyn (Chou et. al, 2010). On GLICOs, a fixed-dose with 4-group ex vivo combination analyses were followed over a period of 1 week and CI values were calculated using invivoSyn (Mao et. al, 2023).

#### Statistical analysis

Data were obtained from three independent experiments and evaluated for their statistical significance with the two-tail unpaired Student’s t test for two groups or analysis of variance (ANOVA) with post hoc corrections in the case of three or more groups. For growth over time, a two-way ANOVA followed by post hoc comparison was employed to analyze and compare the curves. Values were considered statistically significant at *P* below 0.05.

### KEY RESOURCES TABLE

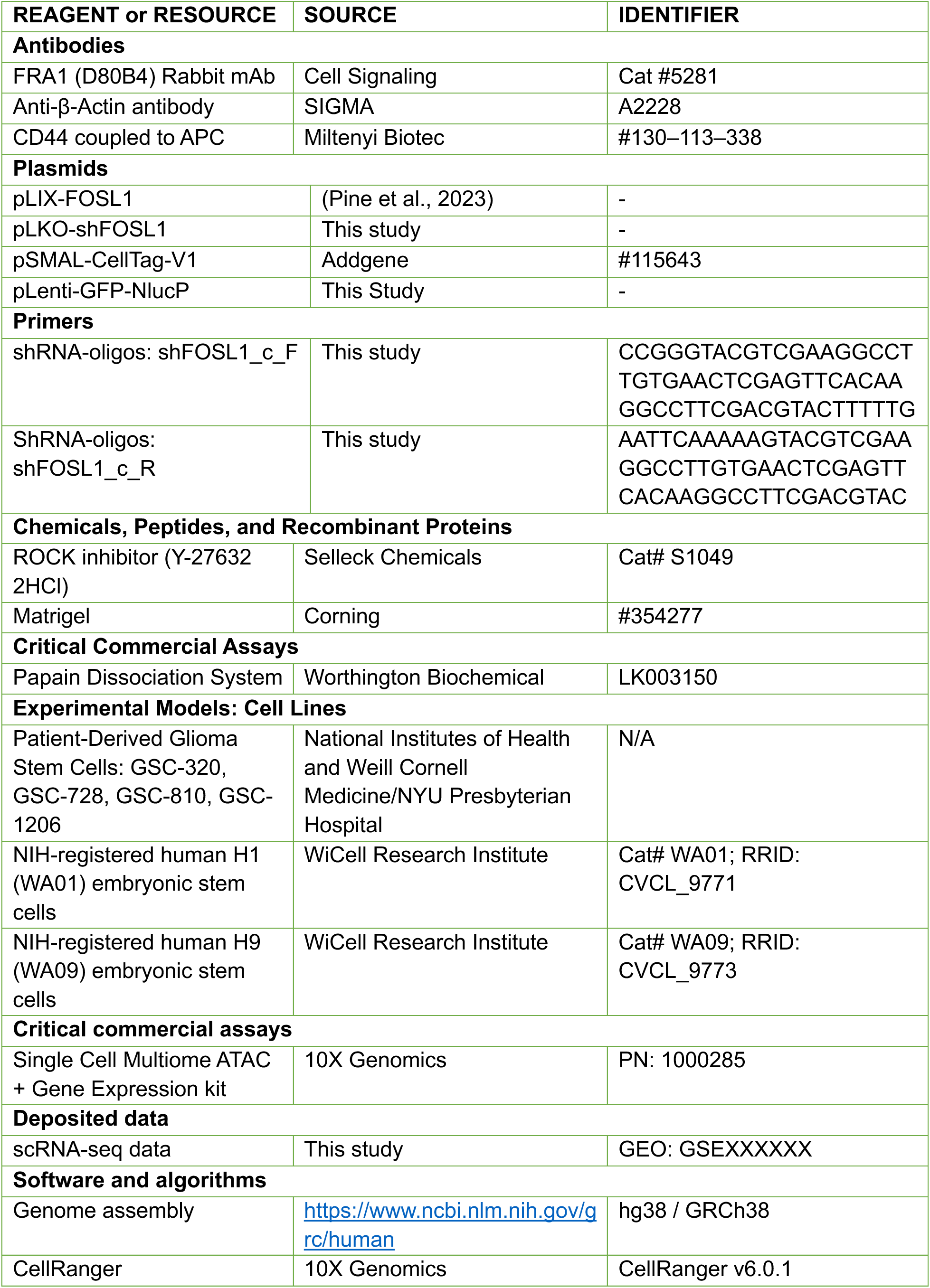
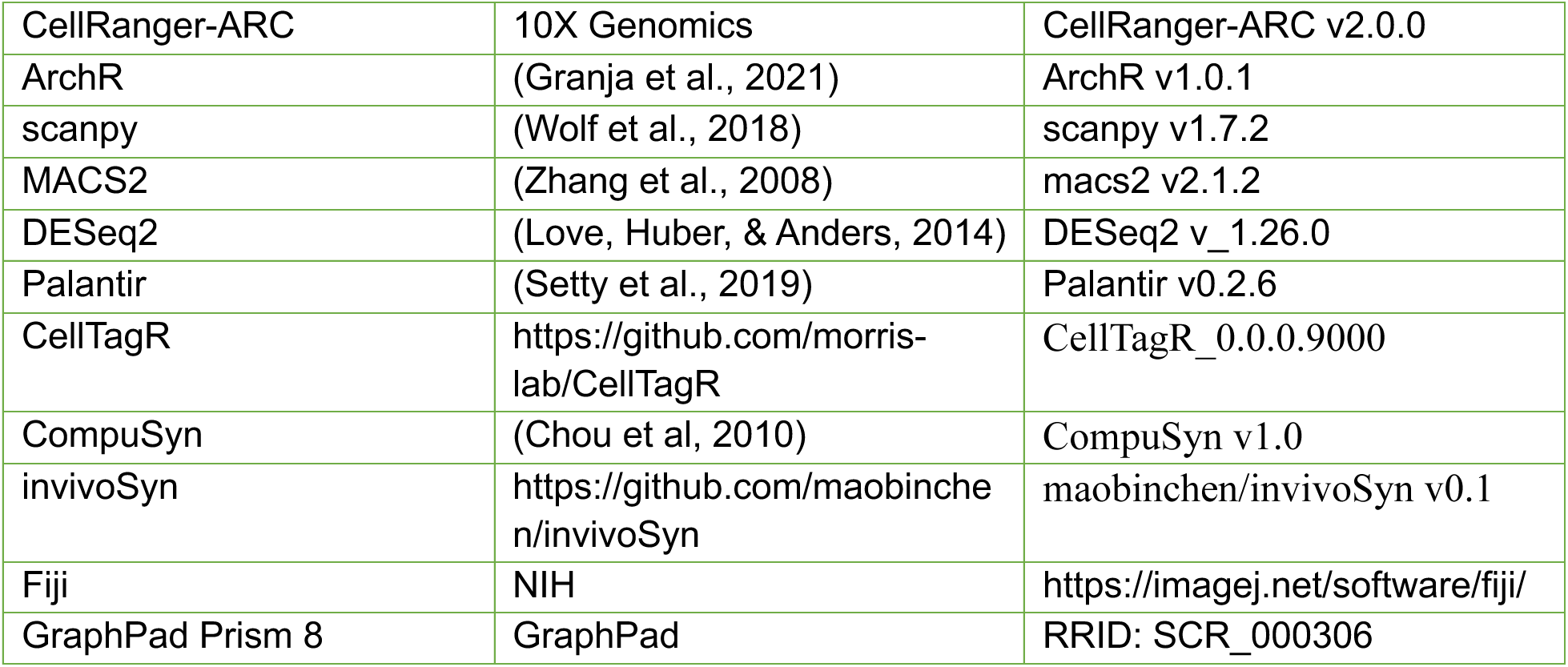

## References

Neftel, C. et al. An integrative model of cellular states, plasticity, and genetics for glioblastoma. Cell 178, 835–849.e21 (2019).

Nicholson, J.G. & Fine, H.A. Diffuse glioma heterogeneity and its therapeutic implications. Cancer Discov. 11, 575–590 (2021).

Linkous, A. et al. Modeling patient-derived glioblastoma with cerebral organoids. Cell Rep. 26, 3203–3211.e5 (2019).

Liu, G. et al. Analysis of gene expression and chemoresistance of CD133+ cancer stem cells in glioblastoma. Mol. Cancer 5, 67 (2006).

Guo, C. et al. CellTag Indexing: genetic barcode-based sample multiplexing for single-cell genomics. Genome Biol 20, 90 (2019).

Pine, A. R. et al. Tumor microenvironment is critical for the maintenance of cellular states found in primary glioblastomas. Cancer Discov. 10, 964–979 (2020).

Chen, Z. et al. FOSL1 promotes proneural-to-mesenchymal transition of glioblastoma stem cells via UBC9/CYLD/NF-κB axis. Mol Ther. 30, 2568–2583 (2022).

Marques, C., et al. NF1 regulates mesenchymal glioblastoma plasticity and aggressiveness through the AP-1 transcription factor FOSL1 eLife 10, e64846 (2021).

Barthel, F. P. et al. Longitudinal molecular trajectories of diffuse glioma in adults. Nature 576, 112–120 (2019).

Suvà, M.L. & Tirosh, I. The Glioma Stem Cell Model in the Era of Single-Cell Genomics. Cancer Cell 37, 630–636 (2020)

Garnier, D., et al. Glioblastoma Stem-Like Cells, Metabolic Strategy to Kill a Challenging Target. Front Oncol. 6, 118 (2019).

Wang, Z. et al. The adaptive transition of glioblastoma stem cells and its implications on treatments. Sig. Transduct. Target. Ther. 6, 124 (2021).

Wang, Y., et al. Direct Pacbio sequencing methods and applications for different types of DNA sequences. 2023.12.12.571020 Preprint at 10.1101/2023.12.12.571020 (2023).

Mao, B. & Guo, S. Statistical Assessment of Drug Synergy from In Vivo Combination Studies Using Mouse Tumor Models. Cancer Research Communications 3, 2146–2157 (2023).

Lu, C. et al. Combined targeting of glioblastoma stem cells of different cellular states disrupts malignant progression. Nat Commun 16, 2974 (2025).

Chou T-C. Drug combination studies and their synergy quantification using the Chou-Talalay method. Cancer Res. 70:440–6 (2010).

Park, J. H. et al. Gene regulatory network topology governs resistance and treatment escape in glioma stem-like cells. Sci Adv 10, eadj7706 (2024).

Antonica, F. et al. A slow-cycling/quiescent cells subpopulation is involved in glioma invasiveness. Nat Commun 13, 1–15 (2022).

Singh, S. et al. Glioblastoma at the crossroads: current understanding and future therapeutic horizons. Sig Transduct Target Ther 10, 1–43 (2025).

Chen, Z. et al. FOSL1 promotes proneural-to-mesenchymal transition of glioblastoma stem cells via UBC9/CYLD/NF-κB axis. Mol Ther 30, 2568–2583 (2022).

Greenwald, A. C. et al. Integrative spatial analysis reveals a multi-layered organization of glioblastoma. Cell 187, 2485–2501.e26 (2024).

